# Characterizing uncertainty in predictions of genomic sequence-to-activity models

**DOI:** 10.1101/2023.12.21.572730

**Authors:** Ayesha Bajwa, Ruchir Rastogi, Pooja Kathail, Richard W. Shuai, Nilah M. Ioannidis

**Author notes:** Emails: {, }. Authors contributed equally to paper (Ayesha Bajwa, Ruchir Rastogi, Pooja Kathail). Co-authorship order was randomly chosen and happens to correspond to love for carrot cake in descending order.

## Abstract

Genomic sequence-to-activity models are increasingly utilized to understand gene regulatory syntax and probe the functional consequences of regulatory variation. Current models make accurate predictions of relative activity levels across the human reference genome, but their performance is more limited for predicting the effects of genetic variants, such as explaining gene expression variation across individuals. To better understand the causes of these shortcomings, we examine the uncertainty in predictions of genomic sequence-to-activity models using an ensemble of Basenji2 model replicates. We characterize prediction consistency on four types of sequences: reference genome sequences, reference genome sequences perturbed with TF motifs, eQTLs, and personal genome sequences. We observe that models tend to make high-confidence predictions on reference sequences, even when incorrect, and low-confidence predictions on sequences with variants. For eQTLs and personal genome sequences, we find that model replicates make inconsistent predictions in >50% of cases. Our findings suggest strategies to improve performance of these models.

## Introduction

Genomic sequence-to-activity models predict molecular phenotypes, such as DNA accessibility, transcription factor (TF) binding, histone modifications, and gene expression, directly from DNA sequence. Numerous deep learning models have been trained for these tasks using experimental assay data from a diversity of cell and tissue types collected by consortia such as ENCODE and Roadmap [1–6]. These models vary significantly in their architectures and the sequence context lengths they consider. Nearly all are trained on human reference genome sequences, which lack the variation present in the human population, but still aim to capture causal effects of regulatory variation. Recent methods show strong performance in predicting expression in massively parallel reporter assays [5, 7] and enhancing functionally informed fine mapping of expression quantitative trait loci (eQTLs) [8]. However, some studies have identified shortcomings of these models, including difficulties in capturing distal regulatory information [9] and inability to consistently predict expression variation across individuals based on their genetic sequence differences [10, 11].

Previous efforts to understand the limitations of these models have focused on evaluating them on biological benchmarks. In this paper, we take a complementary approach of estimating the prediction uncertainty of these models, which can help identify the source of model failures. High-confidence (low uncertainty) incorrect predictions suggest issues such as systematic biases or experimental noise in the training data [12]. Conversely, low-confidence (high uncertainty) predictions, even when correct, could indicate that the training data are insufficient to generalize to unseen test data. For instance, overparameterized models trained on the same data but with different parameter initializations can reach different local minima in the nonconvex loss surface [13, 14]. These models may have similar losses on the training set and even test set, but have different predictions on out-of-distribution inputs, reflecting the underlying uncertainty [15]. Since a primary goal of genomic sequence-to-activity models is to predict the functional effects of regulatory variation, which can viewed as extrapolating to novel sequences not seen at training, distinguishing between these two failure modes can help guide the design of future models. In addition, uncertainty estimation can be used to determine whether high-confidence predictions on some sequences can be trusted, even if the model on average performs poorly. Finally, uncertainty estimation does not require high-quality ground-truth biological data.

Here, we estimate the predictive uncertainty of Basenji2 by evaluating the consistency in predictions made by multiple replicates of the model, trained with different random seeds. Basenji2 makes accessibility, TF binding, histone modification, and gene expression predictions in 128 bp windows using a 131,072 bp sequence as input and ∼55kb receptive field. We use the Basenji2 architecture as it is representative of state-of-the-art genomic sequence-to-activity models and is amenable to training multiple model replicates. We characterize prediction uncertainty across model replicates on four types of sequences: reference sequences, reference sequences perturbed with known TF motifs, eQTLs, and personal genome sequences. We broadly observe that models tend to make high-confidence predictions on reference sequences, even when incorrect, and make low-confidence predictions on sequences with variants.

## Results

### Uncertainty quantification

In a supervised learning setup, with inputs *x* and outputs *y* related through the joint distribution *q*(*x, y*), we can train a model *p*(*y* | *x, θ*) on a finite dataset 𝒟 = {(*x*_1_, *y*_1_), …, (*x*_*n*_, *y*_*n*_)} ∼ *q*(*x, y*) with *n* training examples. Adopting a Bayesian framework, we can decompose the predictive uncertainty into data (aleatoric) and model (epistemic) uncertainty [16, 17]:

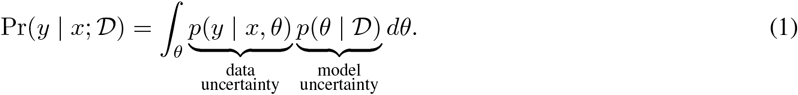

Data uncertainty refers to irreducible (does not decrease as *n* increases) uncertainty in the complexity of the data, for example due to class overlap or measurement noise. On the other hand, model uncertainty refers to reducible (decreases as *n* increases) uncertainty in estimating the true parameters *θ*^*^ given a finite dataset 𝒟. Previous methods proposed to approximate the integral in Eq. 1 include variational Bayesian neural networks [18] and Monte-Carlo dropout [19]. However, a surprisingly performant and effective approach to estimate predictive uncertainty, even under dataset shift, is to train a deep ensemble of *M* models 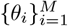 that differ only in their random seeds [20, 14, 21]:

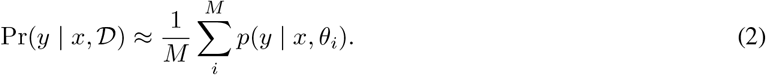

For Basenji2 [4], the conditional *p*(*y* | *x, θ*_*i*_) is the likelihood of *y* under a Poisson distribution whose mean and variance equal the model output, 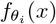. Therefore, Eq. 2 is a convex combination of independent Poisson random variables and defines a Poisson mixture model (a Poisson mixture model is always overdispersed and therefore not Poisson itself [22]). In practice, we typically aim to estimate uncertainty on binary classification tasks that involve complicated transformations of model predictions. For example, for eQTL sign prediction, we would like to estimate the probability that the eQTL increases expression: Pr(*y*_alt_ − *y*_ref_ > 0 | *x*_alt_, *x*_ref_; 𝒟). Unfortunately, there is no computationally tractable method to compute the probability mass function for the difference of two Poisson mixture models. Accordingly, we approximate this probability by determining the fraction of models in the ensemble whose point estimates suggest increased expression:

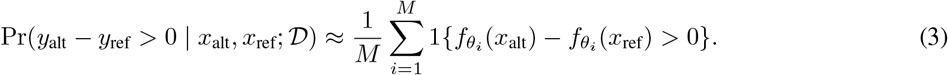

The approximation in Eq. 3 is coarse. However, since our goal is not to produce rigorous uncertainty estimates but instead to understand the types of sequences for which these models have high predictive uncertainty, it suffices for our analysis. When the probability in Eq. 3 is equal to 0 or 1 (i.e. all replicates predict the same direction change), we say that the replicates are *consistent*. We further breakdown the consistent category into *consistently correct* and *consistently incorrect* when ground truth data is available. When the probability is in (0, 1) (i.e. the replicates disagree in their predicted direction change), we say that the replicates are *inconsistent*.

### Ensemble training

We train 5 Basenji2 replicates using only the human training data from the Basenji2 dataset [4] and evaluate consistency in the predictions of the replicates. All replicates were trained with the same data and hyperparameters (learning rate, batch size, early stopping, etc.), differing only in their random seeds. Specifically, the replicates differed only in their random parameter initialization, the random sampling of training examples during mini-batch optimization, and random dropping of neurons during training due to dropout. All five replicates have similar performance on the fixed test split of the Basenji2 dataset (Fig. S1). We hypothesize that the gap between the replicates and the Basenji2 model in [4] is the lack of multi-species training (the original also trains on mouse data). While recent works have ensembled replicate models to improve prediction performance [6, 23], to our knowledge, we are the first to train replicate models to assess prediction uncertainty of genomic sequence-to-activity models.

### Reference genome predictions are largely consistent across models, even when incorrect

We first assess the consistency between replicates on reference genome sequences held out during training. For each of the 5,313 Basenji2 tracks, we compute the Pearson correlation between predictions from replicates 1 and 2 on held out reference sequences (Fig. 1a). We observe high correlation between replicates (median Pearson’s *r >* 0.9) for all assays, suggesting that the replicates agree on relative activity differences across the reference genome. Predictions for CAGE tracks are slightly less correlated between replicates than predictions for other assays (DNase-seq/ATAC-seq, TF ChIP-seq, and histone modification ChIP-seq). Previous studies have noted that consistency in predictions does not entail consistency in input feature attributions [24, 25], which reflect the relative importance of input features to a prediction. However, gradient saliency maps for CAGE predictions (in the GM12878 cell line) for genes unseen during training show that replicates tend to agree on the importance of regulatory regions (Fig. 1b).

**Figure 1:**
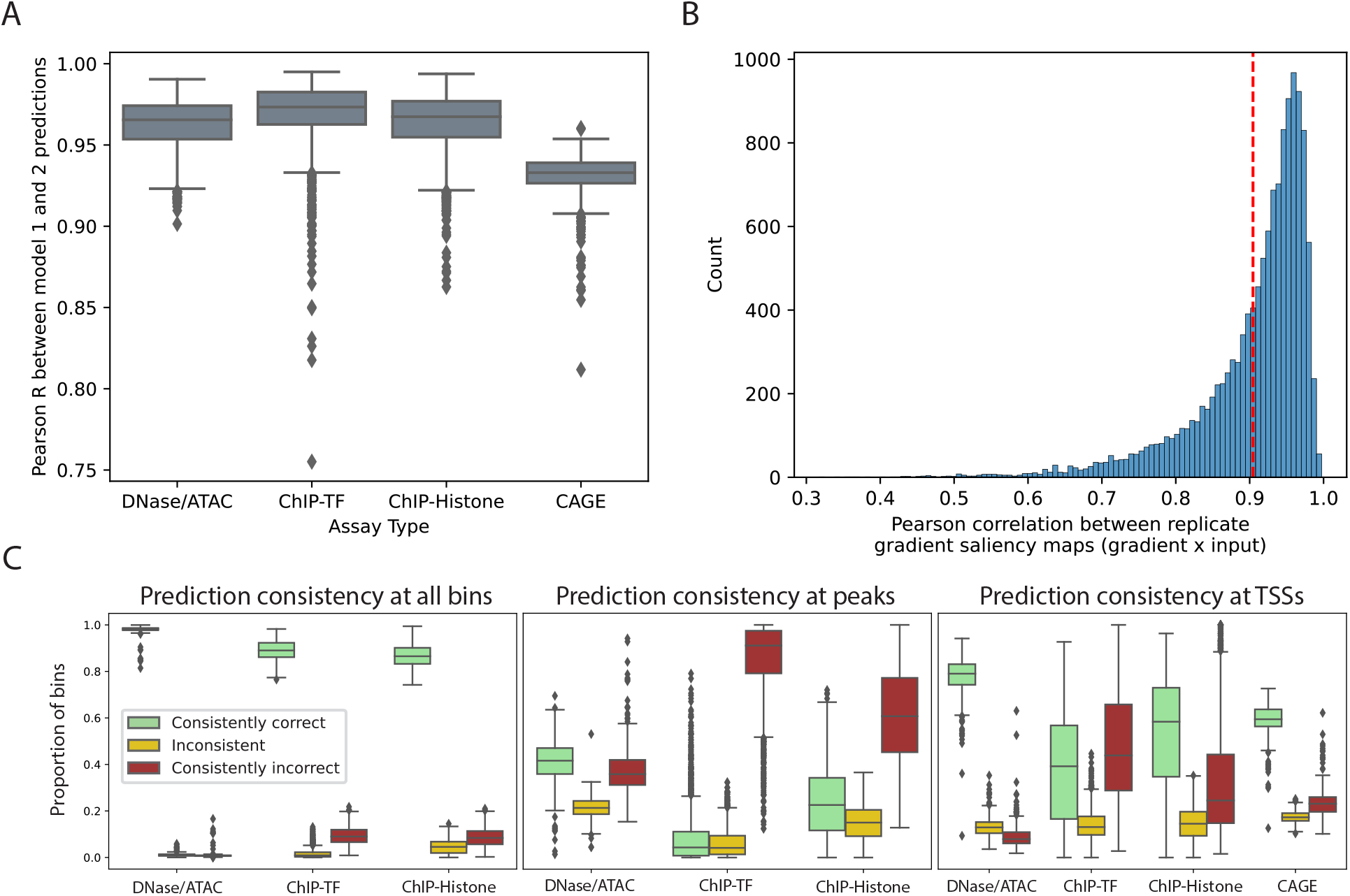
Reference genome predictions are largely consistent across replicates, even when incorrect. (a) For each of the 5,313 Basenji2 tracks, we report the Pearson correlation between predictions from replicate models 1 and 2 on reference genome sequences held out during training. Reference genome predictions are highly correlated across the two replicates, and predictions for CAGE tracks are less consistent than for other assays. (b) For genes held out during training, we display the distribution of Pearson correlations between gradient saliency maps from all pairs of replicates. The red dashed line indicates the mean Pearson correlation. (c) We report the proportion of test sequences that are classified consistently correctly, inconsistently, or consistently incorrectly in a binary peak prediction task. We distinguish performance on all bins (left), peaks (center), and peaks at transcription start sites (right).

Next, we ask whether consistently predicted reference sequences are more often correct with respect to the experimental measurements. We binarize the experimental and predicted activity levels for held out reference sequences by calling peaks separately per track (Methods). Using these binary peak calls, we classify reference sequences into one of three categories: consistently correct (all 5 replicates agree with the experimental peak label), inconsistent (3 or 4 of the replicates agree with each other), or consistently incorrect (all 5 replicates disagree with the experimental peak label). Most reference sequences are classified consistently correctly (median proportion > 0.8 for all assays), and only a small fraction of all sequences are predicted inconsistently (median proportion < 0.1 for all assays) (Fig. 1c, left). However, within the much smaller subset of sequences corresponding to experimental peaks, approximately 20% are predicted inconsistently across replicates (Fig. 1c, center). Strikingly, for peak sequences predicted consistently by all five replicates, a large fraction are consistently incorrect. We acknowledge that reliance on a peak calling threshold is a potential shortcoming, and that the choice of threshold may influence the proportion of sequences in each category.

Lastly, we subset further to sequences containing transcription start sites (TSSs) that are also experimentally determined peaks. On average, across CAGE tracks, ∼60% of these sequences are predicted consistently correctly, ∼20% are predicted consistently incorrectly, with the remaining ∼20% predicted inconsistently (Fig. 1c, right). In comparison to our analysis of all peaks (Fig. 1c, center), we observe a similar proportion of inconsistently predicted sequences (∼20%). However, for consistent predictions, TSS peaks are predicted correctly much more often than non-TSS peaks.

To explore potential systematic differences in the types of sequences predicted consistently or not, we analyze the predictions for two epigenetic tracks (DNase-seq and CTCF ChIP-seq) in GM12878 cells. We evaluate whether peak sequences in each of the three consistency categories differ on five different attributes: GC content, TSS distance, evolutionary conservation (phyloP), experimentally measured activity level (target), and mean predicted activity level across replicates (Fig. S2). Peak sequences predicted consistently incorrect have significantly lower experimentally measured activity levels. Further, consistently correctly predicted peak sequences show more resemblance to promoters: they are more proximal to the TSS, have higher GC content, and are more evolutionarily conserved. This is consistent with previous reports that current sequence-to-activity models capture gene expression determinants in promoters, but struggle with distal sequences [9]. On all five attributes, inconsistently predicted peak sequences display characteristics in between those of the consistently correctly and consistently incorrectly predicted peak sequences.

### Replicates disagree more on the effect of mutations in TF motifs than on the effect of canonical TF motifs

Deep learning models can learn gene regulatory syntax in part by learning TF motifs in their first layer convolutional filters [26]. Differences in the learned effects of TF motifs may contribute to inconsistent predictions across replicates. To test this hypothesis, we compute TF activity scores (the difference in predicted activity at the TSS for motif-inserted versus endogenous background sequences) for all human CIS-BP motifs using each replicate at four different fixed positions upstream of each gene’s TSS (Fig. 2a & Methods). We focus on the TF activity score sign, reasoning that replicates may have slight differences in TF activity magnitude but should agree on whether a TF increases or decreases activity if they have learned similar gene regulatory syntax. For each prediction track, we compute the fraction of TFs with inconsistently predicted directional effects across replicates (Fig. 2b, light gray). We observe differences in the consistency of predicted TF activity signs for different assays. When motifs are inserted very proximal (10bp upstream) of the gene’s TSS, we observe that TF activity scores are most consistent for CAGE tracks compared to other assays. We also find that–in general–replicate consistency decreases as the TF is inserted farther from the gene’s TSS, although consistency for DNase and ATAC tracks is relatively stable across different insertion positions. We note that, compared to TSS-proximal regulatory elements, distal regulatory elements tend to have smaller effects on TSS activity, and current models underpredict the effect of distal regulatory elements on gene expression, both of which may contribute to greater inconsistency for distal TF motif insertions [9].

**Figure 2:**
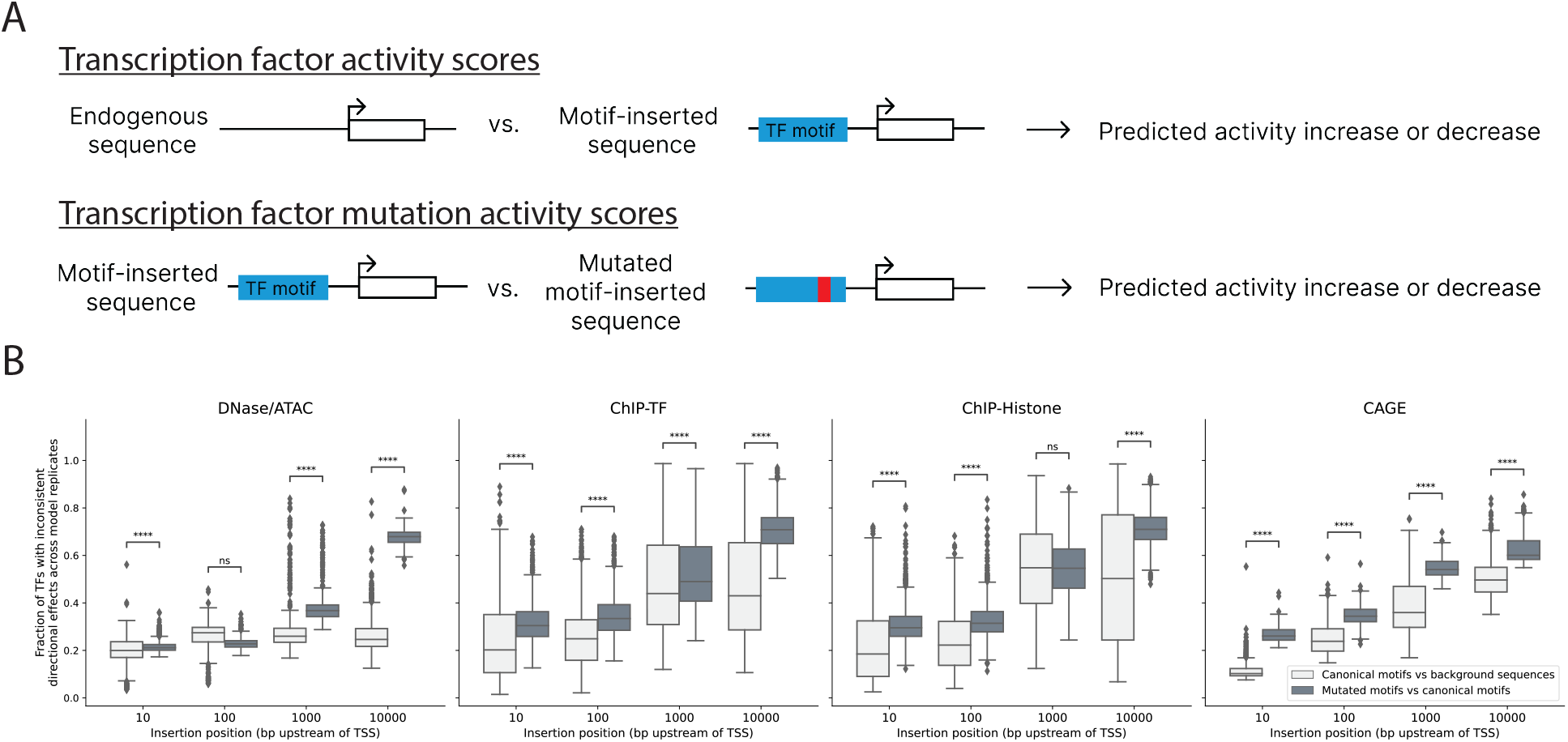
Replicates display greater inconsistency at predicting effects of mutations in TF motifs compared to canonical TF motifs. (a) Graphical depiction of the TF activity score and TF mutation activity score calculations. TF activity scores were calculated by comparing model predictions for endogenous background sequences versus background sequences with a canonical TF motif inserted at a fixed location upstream of a gene’s TSS. TF mutation activity scores were calculated by comparing model predictions for canonical motif-inserted sequences versus sequences where a single base-pair mutation was made to the canonical motif. Scores were averaged over 100 different background sequences. (b) We compute TF activity scores and TF mutation activity scores by inserting motifs at four different fixed positions–10bp, 100bp, 1000bp, and 10,000bp–upstream of each selected gene’s TSS. For all Basenji2 prediction tracks, we report the fraction of TFs with inconsistent predicted directional effects across model replicates for both TF activity scores (canonical motifs versus background sequences) and TF mutation activity scores (mutated motifs versus canonical motifs). In all but two cases, we observe greater inconsistency in TF mutation activity scores than TF activity scores (one-sided Mann-Whitney U test, **** indicates a Benjamini-Hochberg corrected p-value < 1e-4)

Since genomic sequence-to-activity models can be used to understand the functional effects of regulatory variation, we consider the effect of mutations in TF motifs. We hypothesized that it may be easier for replicates to learn consistent effects for a canonical TF motif compared to a single base pair mutation in a motif. For each human CIS-BP TF motif, we select a mutation likely to disrupt activity (by mutating the lowest entropy PWM position to its lowest probability base) and compute TF mutation activity scores for insertions at the same four positions upstream of each gene’s TSS (Fig. 2a & Methods). For each track, we report the fraction of TF mutations with inconsistent predicted directional effects across replicates (Fig. 2b, dark gray). For almost all assays and distances, there is greater inconsistency in predicting the effect of a mutation to a TF motif compared to predicting the effect of the canonical motif. The trend is more pronounced for perturbations farther from the TSS. The correlation between consistency of TF mutation activity scores (calculated using perturbations 10bp upstream of each gene’s TSS) and mutation probabilities (based on PWMs) shows that higher probability mutations, which are less likely to disrupt TF binding, have less consistent predictions (Fig. S3). Therefore, by selecting a strongly disruptive (low probability) mutation to each motif, our analysis may represent a lower bound on inconsistency in mutation prediction, as less disruptive mutations are likely to have more inconsistent predictions.

### eQTL sign predictions show high replicate inconsistency

To further quantify uncertainty in variant effect predictions, we utilize a dataset of fine-mapped expression quantitive trait loci (eQTLs) from the Genotype-Tissue Expression Consortium (GTEx) [27] and matched negative variants [5]. We select thirteen tissues with matched DNase-seq and CAGE tracks among the 5,313 Basenji2 tracks. Using each replicate, we compute variant effect predictions (SAD scores) for all fine-mapped eQTLs and matched negatives (Methods). Fine-mapped eQTL predictions show higher pairwise correlations between replicates than matched negative predictions (Fig. S4), suggesting that replicates make more consistent predictions for putatively causal variants.

We focus on replicate agreement in predicting the directional effect of fine-mapped eQTLs (whether the eQTL alternate allele increases or decreases gene expression compared to the reference). Using predicted SAD signs from matching CAGE tracks, 55% of eQTLs (pooled across tissues) have inconsistent predictions (about 50-60% in individual tissues, Fig. S5), 29% have consistently correct predictions, and 16% have consistently incorrect predictions (Fig. 3a). Of the consistently predicted eQTLs, about 65% are correct. Predictions from DNase-seq tracks are inconsistent for about 40-50% of eQTLs (Fig. S6a, d) and, similar to CAGE, about 65% of consistent DNase predictions are correct.

**Figure 3:**
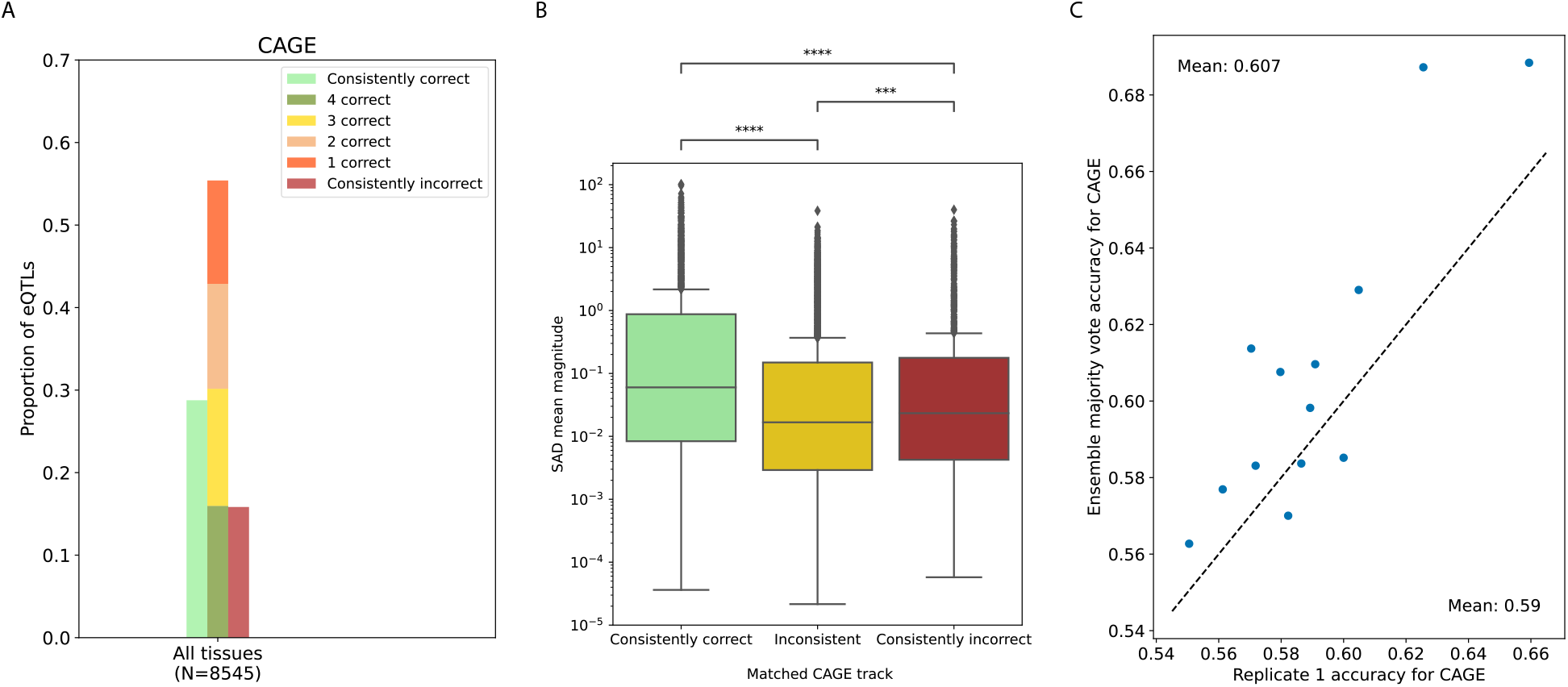
eQTL sign predictions from tissue-matched CAGE tracks have high inconsistency across replicates. (a) We show the proportion of fine-mapped eQTLs with consistently correct, inconsistent, and consistently incorrect eQTL effect sign predictions across replicates. About 55% of eQTLs have inconsistent predictions, while 29% are consistently correct and 16% are consistently incorrect. (b) eQTLs with inconsistent sign predictions tend to have smaller predicted effect sizes (mean of SAD score magnitude across replicates) in the tissue-matched CAGE track. (c) A comparison of accuracy for eQTL sign prediction shows that the ensemble majority vote does not substantially or consistently outperform a single replicate. Each point represents the fine-mapped eQTL set of a different tissue.

We next investigate whether there are systematic differences between eQTLs with consistent and inconsistent sign predictions. Predicted effect sizes, computed as the mean SAD magnitude across replicates, tend to be larger for eQTLs with consistent predictions when using either the tissue-matched CAGE (Fig. 3b) or DNase (Fig. S6b) tracks, suggesting small predicted differences between alleles are less likely to be consistent. Additionally, we check if eQTLs with inconsistent sign predictions are overrepresented in certain regulatory regions, using their tissue-matched chromHMM annotation (Fig. S7) [28, 29]. However, we find that at least 40% of eQTLs have inconsistent sign predictions in all annotations, irrespective of the annotation’s distance to TSS or whether the annotation corresponds to an active or repressed state.

We then compare the performance of a single replicate to an ensemble majority vote on sign prediction. The ensemble majority vote marginally outperforms a single replicate in 10/13 tissues for tissue-matched CAGE tracks (Fig. 3c) and in 5/13 tissues for tissue-matched DNase tracks (Fig. S6c). For the six tissues with the most fine-mapped eQTLs, the ensemble often provides a slight improvement for TSS-proximal eQTLs (Fig. S8), consistent with previous results [5].

### Replicates vary substantially in predictions on personal genomes

Recent work [11, 10] has found that current genomic sequence-to-activity models are unable to explain variation in gene expression across individuals based on their personal genome sequences. Huang et al. [10] evaluated four models–Xpresso, ExPecto, Basenji2, and Enformer–using paired whole-genome sequencing and RNA-seq data from lymphoblastoid cell lines (LCLs) of 421 individuals in the Geuvadis Consortium, and found that the models strongly disagree with one another on the predicted direction of genetic effects on expression. Motivated by this observation, we seek to determine whether disagreement between model classes is driven by predictive uncertainty.

We made predictions with the five replicates on 3259 genes with significant eQTLs in the Geuvadis dataset (Methods). There is significant disagreement in replicate predictions (Fig. 4a): in 2181 (67%) genes, replicates differ in the signs of their cross-individual correlations (Spearman correlation between predicted expression and measured RNA-seq across individuals). Comparing the cross-individual correlations of two replicates, we see that many genes have similar correlation magnitudes even if they have opposite signs, as observed by enrichment along the *y* = −*x* diagonal (Fig. 4b). This matches the finding in Huang et al. [10] that different model classes such as Basenji2 and Enformer often make predictions with more consistency in the magnitudes of their cross-individual correlations than in their signs. Our analysis suggests this phenomenon is likely not primarily a result of differences in model architecture or training procedure, but rather a result of predictive uncertainty. As before, we assess if prediction performance can be improved using an ensemble of the five replicates, but find that it does not substantially outperform a single replicate (Fig. 4c).

**Figure 4:**
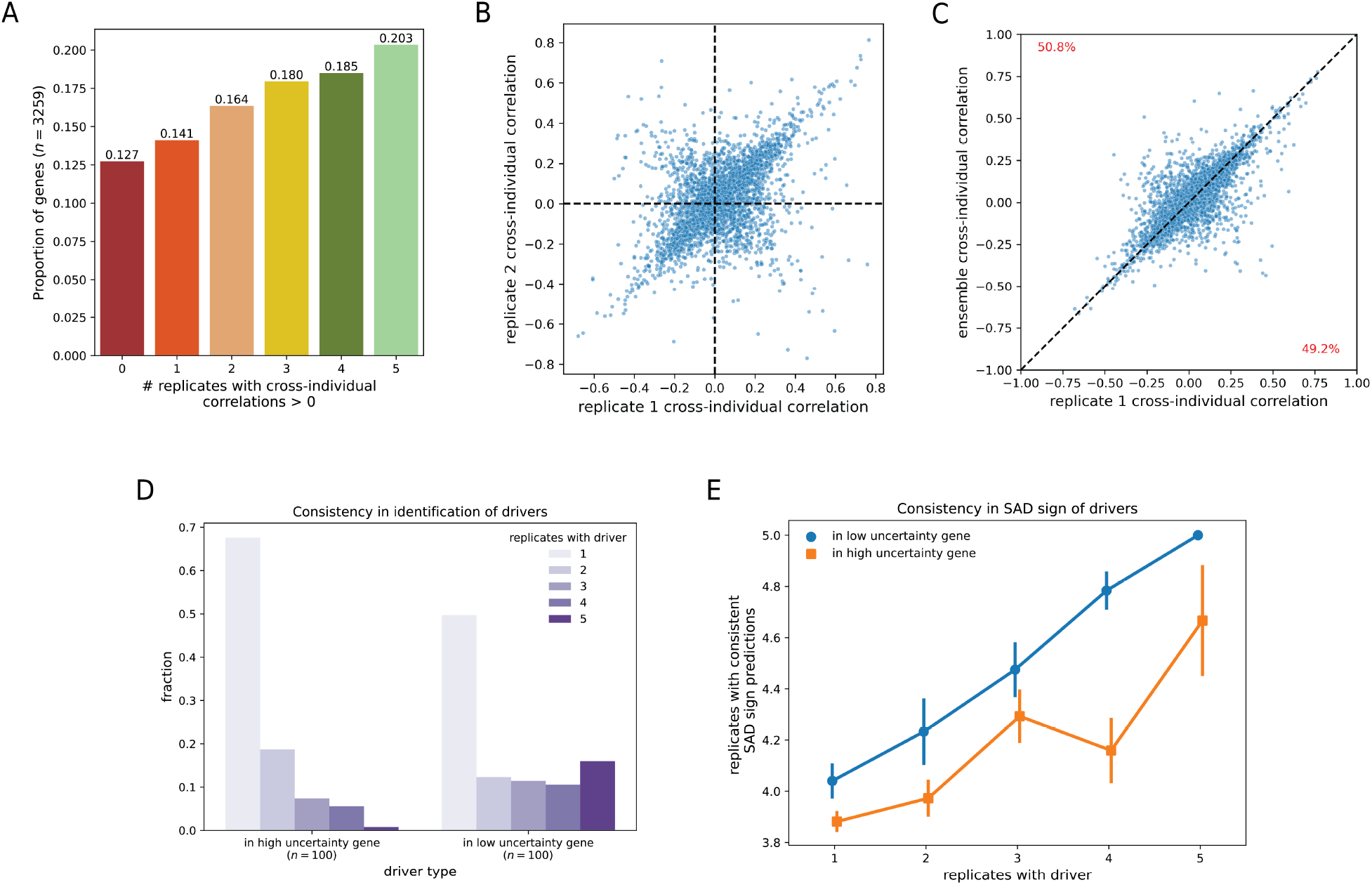
Uncertainty in predictions on personal genome sequences. (a) We segment the 3259 genes analyzed in Huang et al. [10] by the number of replicates with cross-individual correlations greater than 0. For each gene, the cross-individual correlation is calculated as the Spearman correlation between model predictions and measured RNA-seq across individuals. (b) We compare the cross-individual Spearman correlations of replicates 1 and 2. (c) We compare the cross-individual Spearman correlations of replicate 1 and an ensemble model. The ensemble model averages an individual’s predicted expression rank across all five replicates. (d) We show the number of drivers that are identified reproducibly in multiple replicates, stratified by whether these drivers are found in genes with high or low uncertainty. (e) We plot the number of replicates (3-5) that agree on the SAD score sign of a driver, stratified both by the number of replicates that identify the driver and if the driver was found in an high or low uncertainty gene.

To better understand the source of predictive uncertainty, we compared 100 high uncertainty genes to 100 low uncertainty genes, chosen based on the variance in the replicate cross-individual correlations. Within this subset of genes, for each replicate, we identified drivers–single nucleotide variants that explain most of the variance in predictions–using the approach described in [11]. Similar to [11], we identify a small number (1-6) of drivers per gene. Drivers in high uncertainty genes tend to be unique to a single replicate, whereas drivers in low uncertainty genes are often found in all five (Fig. 4d). Further, we find that replicates disagree more often on the directional effect of drivers in high uncertainty genes than in low uncertainty genes, even when conditioning on the number of replicates that identify a variant as a driver (Fig. 4e). Cumulatively, these observations suggest significant uncertainty not only in determining the directional impact of a variant on gene expression but also in identifying the variants that affect expression.

## Conclusion

We analyze uncertainty in the predictions of genomic sequence-to-activity models by measuring prediction consistency across five replicate Basenji2 models, when applied to reference genome sequences, reference genome sequences perturbed with TF motifs, eQTLs, and personal genome sequences. For held out reference sequences, which are similar in distribution to the training data, predictions show high replicate consistency. However, for sequences that require models to generalize to out-of-distribution regulatory variation – eQTLs and personal genome sequences – predictions show high replicate inconsistency. Surprisingly, consistent predictions for both reference and variant sequences are often incorrect.

These results have implications both for the application of current models and the development of future models. In other domains – including biological applications such as protein design – prediction uncertainty estimates have been employed to determine when to trust a model’s predictions [30–32]. As genomic sequence-to-activity models are increasingly applied to variant interpretation and sequence design problems, we believe accurate quantification of prediction uncertainty can be a useful practical tool. Our initial exploration of uncertainty quantification is based on ensemble prediction consistency. Further work is needed to develop rigorous, calibrated uncertainty estimates for genomic sequence-to-activity models; methods which calculate statistically rigorous uncertainty intervals (e.g. conformal prediction), rather than point estimates alone, are a promising direction.

Our characterization of uncertainty also sheds light on the failure modes of current models. The high degree of inconsistency observed for predictions on variant sequences suggests that models are struggling to generalize to these out-of-distribution inputs. Integrating additional sequence diversity into model training, as some works have suggested [33, 11], may help overcome this limitation. For example, our TF motif insertion analysis revealed that replicates make less consistent predictions about the effects of mutations to TF motifs compared to the effects of canonical motifs, suggesting that in vitro TF binding assays (i.e. SELEX) may be a useful source of training data.

## Methods

### Ensemble model training

We train five replicate models using the Basenji2 model architecture, training procedure, and human training data [4]. We use the Basenji Github repository for model training [34].

### Reference sequence predictions

To make predictions for held out reference sequences, we average predictions over the forward and reverse complement sequence and 1-nucleotide sequence shifts to the left and right. This same averaging procedure is used when making predictions in all subsequent analysis. For computational tractability, we downsample the held-out reference sequences ten-fold in all downstream analysis. We binarize experimental and predicted activity levels for held out reference sequences by calling peaks separately per track, using the method in [34]. Briefly, for each experimental and predicted track, we called peaks using a Poisson model parameterized by a *λ* corresponding to the mean activity across all 128bp bins and applied a 0.01 FDR cutoff.

### Gradient saliency maps

For the 1308 protein-coding genes whose TSS is within a Basenji2 test sequence, we compute the gradient of the GM12878 CAGE prediction (averaged over the central ten bins) with respect to the input reference sequence nucleotides. We sum the absolute value of gradients in 128 bp windows to obtain non-negative contribution scores per bin.

### TF motif activity scores

Inspired by the motif insertion approach in Yuan and Kelley [35], we select a set of 100 gene TSSs to use as endogenous background sequences. These 100 genes were held out from the Basenji2 training data and predicted consistently correctly by the replicates across the greatest number of tracks. For each TF in the CIS-BP motif database [36] and each background sequence, we sample a motif sequence from the TF’s PWM and insert it at a fixed position upstream of the gene TSS. For each model and prediction track, we calculate a TF activity score as the difference in predictions for motif-inserted sequences versus background sequences for the central two 128bp bins (because the TSS is at the junction of these bins). We calculate TF activity scores at four different motif insertion positions – 10bp, 100bp, 1000bp, and 10,000bp – upstream of each gene’s TSS, to assess how consistency in TF activity scores varies for proximal versus distal motifs.

We also consider the effect of mutations on TF activity. For each TF, we simulate a mutation to its motif by selecting the lowest entropy position of its PWM and mutating it to the lowest probability base. We calculate TF mutation activity scores as the difference in predictions for sequences containing mutated motifs versus sequences containing canonical motifs (described above) for the central two 128bp bins.

### GTEx eQTLs

We obtain GTEx v8 eQTLs fine-mapped using SuSiE [37, 8] and matched negative variants from the Supplementary Data in Avsec et al. [5]. We filter out variants which have opposite directions of effect on different genes, as well as variants which are further from the TSS than the receptive field of the Basenji2 architecture allows. Then, for each variant, we retain only the gene-variant pair for the closest gene. To compute variant effect predictions for each track, we subtract the reference allele prediction from the alternate allele prediction across the central three 128bp prediction bins centered at the variant to obtain an absolute SAD (SNP activity difference) score.

### Personal genomes

We predict lymphoblastoid cell line (LCL) gene expression for 421 individuals in the Geuvadis consortium with both phased whole-genome sequencing and LCL RNA-seq data. We focus on the 3259 genes with a significant eQTL in the European cis-eQTL analysis. Following the approach detailed in Huang et al. [10], we construct personal haplotype sequences with single nucleotide variants inserted. For each haplotype, we average predictions for the GM12878 LCL CAGE track over the central ten bins surrounding the TSS. We then average haplotype predictions to obtain individual predictions. For each replicate, we identify drivers using the approach described in Sasse et al. [11] for the 100 genes with the highest uncertainty and 100 genes with the lowest uncertainty. We define a gene’s uncertainty by the variance in the cross-individual correlations of the five replicates and only consider genes where the mean of the cross-individual correlation magnitude is at least 0.1.

## Data and code availability

All datasets used in this study are publicly available. Code to train replicates, make model predictions, and generate figures is available at https://github.com/ni-lab/basenji2_uncertainty.

## Acknowledgements

We thank Gabriel Loeb and Ioannidis lab members for helpful discussions. This work was partially supported by the U.S. National Institutes of Health grant R00HG009677, an Okawa Foundation Research Grant, and a grant from the UC Noyce Initiative for Computational Transformation. N.M.I. is a Chan Zuckerberg Biohub San Francisco Investigator.

## A Appendix

### Supplementary Figures

**Figure S1:**
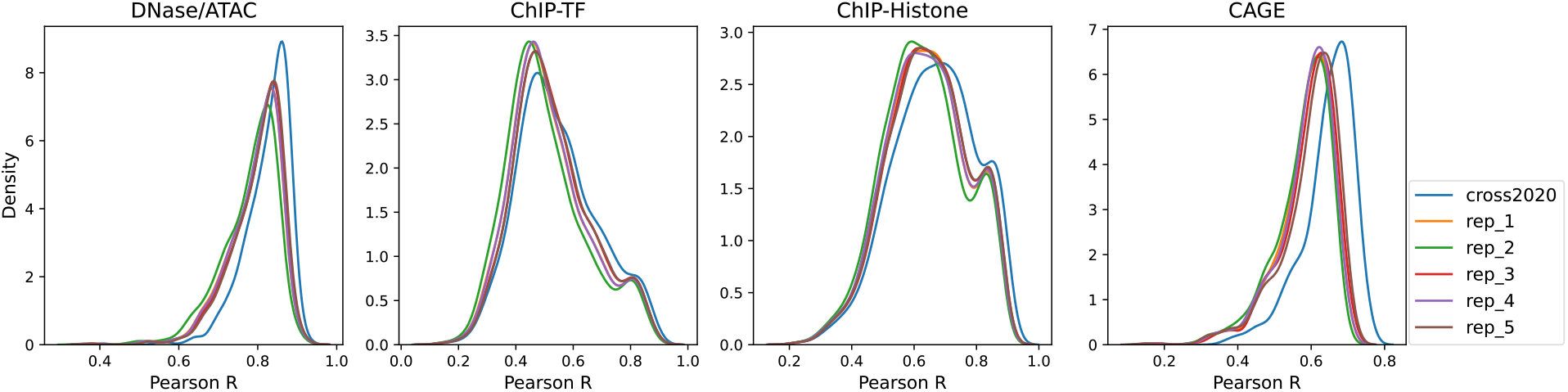
Test set performance of replicates. Pearson correlation on held-out test data across prediction tracks for each assay.

**Figure S2:**
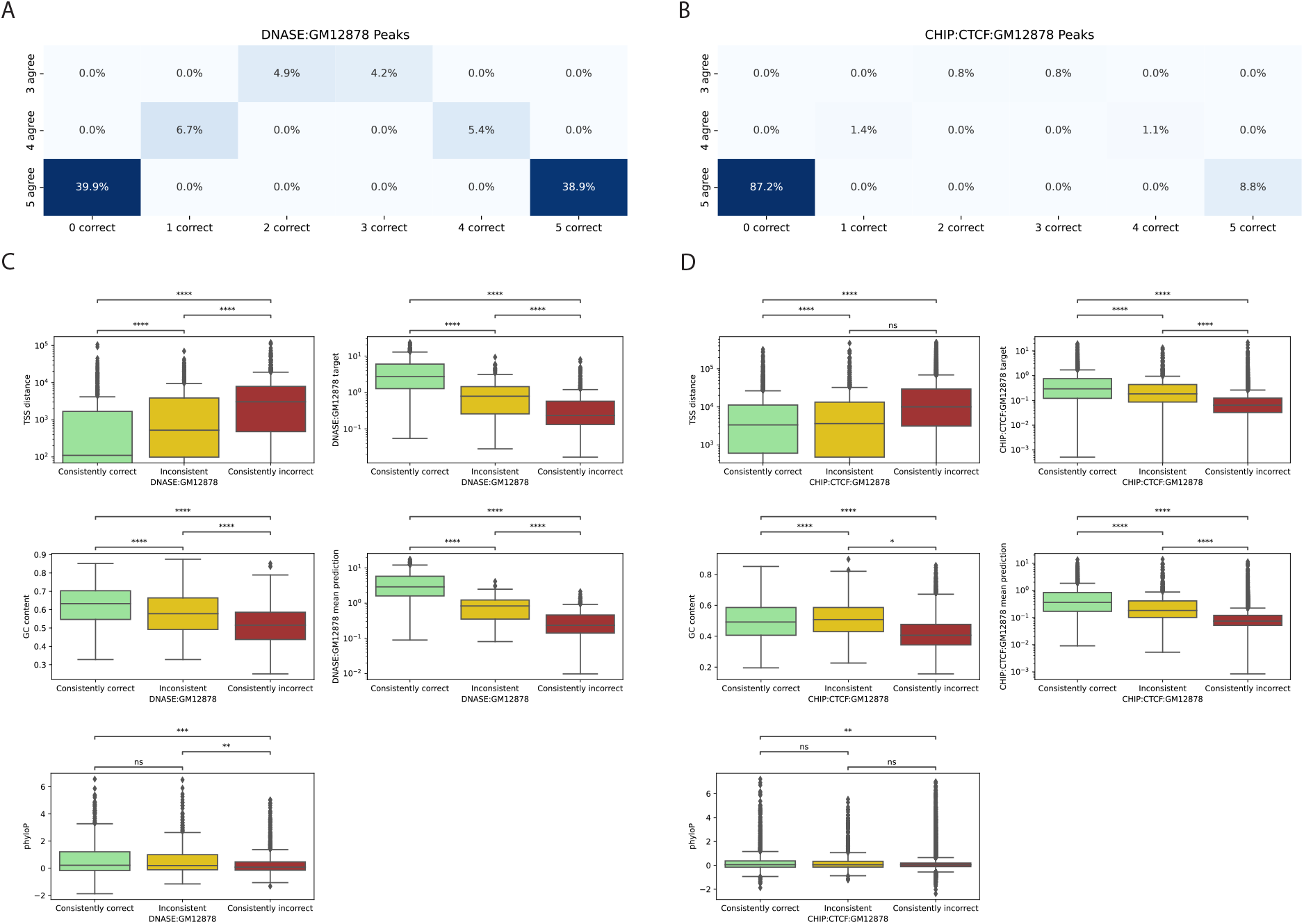
Consistently correctly predicted peaks are gene-proximal, have higher GC content, and are evolutionarily conserved. For (a) DNase-seq and (b) CTCF ChIP-seq in GM12878 cells, we display heatmaps of prediction consistency and correctness across peak sequences. For (c) DNase-seq and (d) CTCF ChIP-seq in GM12878 cells, we measure differences in the sequences falling into each of the three consistency categories (consistently correct vs. inconsistent vs. consistently incorrect) across five attributes – TSS distance, GC content, evolutationary conservation (phyloP), experimentally measured activity level (target) and mean predicted activity level.

**Figure S3:**
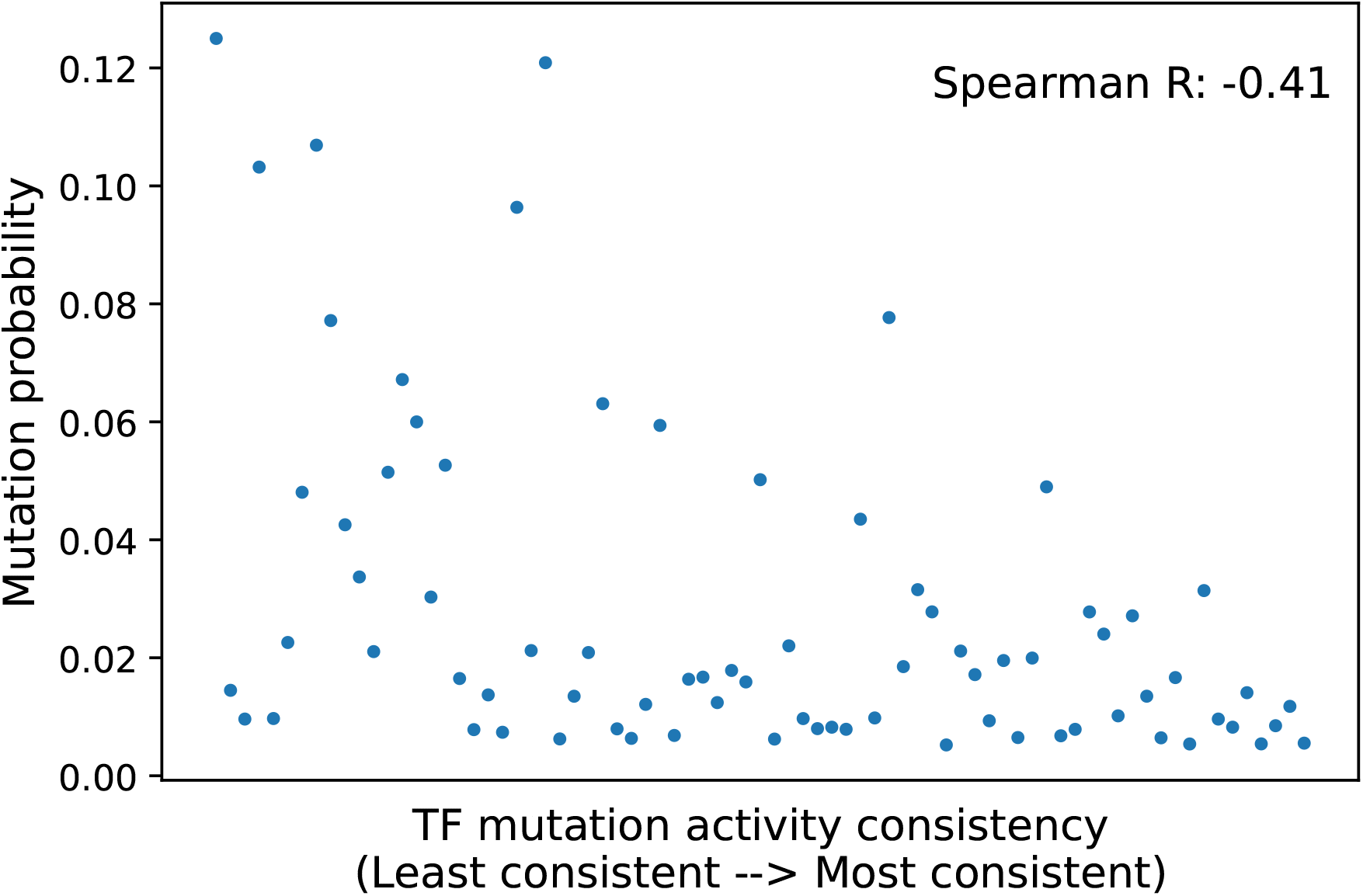
Higher probability (less disruptive) mutations to TF motifs have less consistent predictions across replicates. For tested mutations to TF motifs with probability greater than 0.005 according to the TF’s PWM, we plot consistency in the TF mutation activity scores (calculated using perturbations 10bp upstream of the TSS) across replicates versus mutation probability. We observe that higher probability mutations have less consistent predictions.

**Figure S4:**
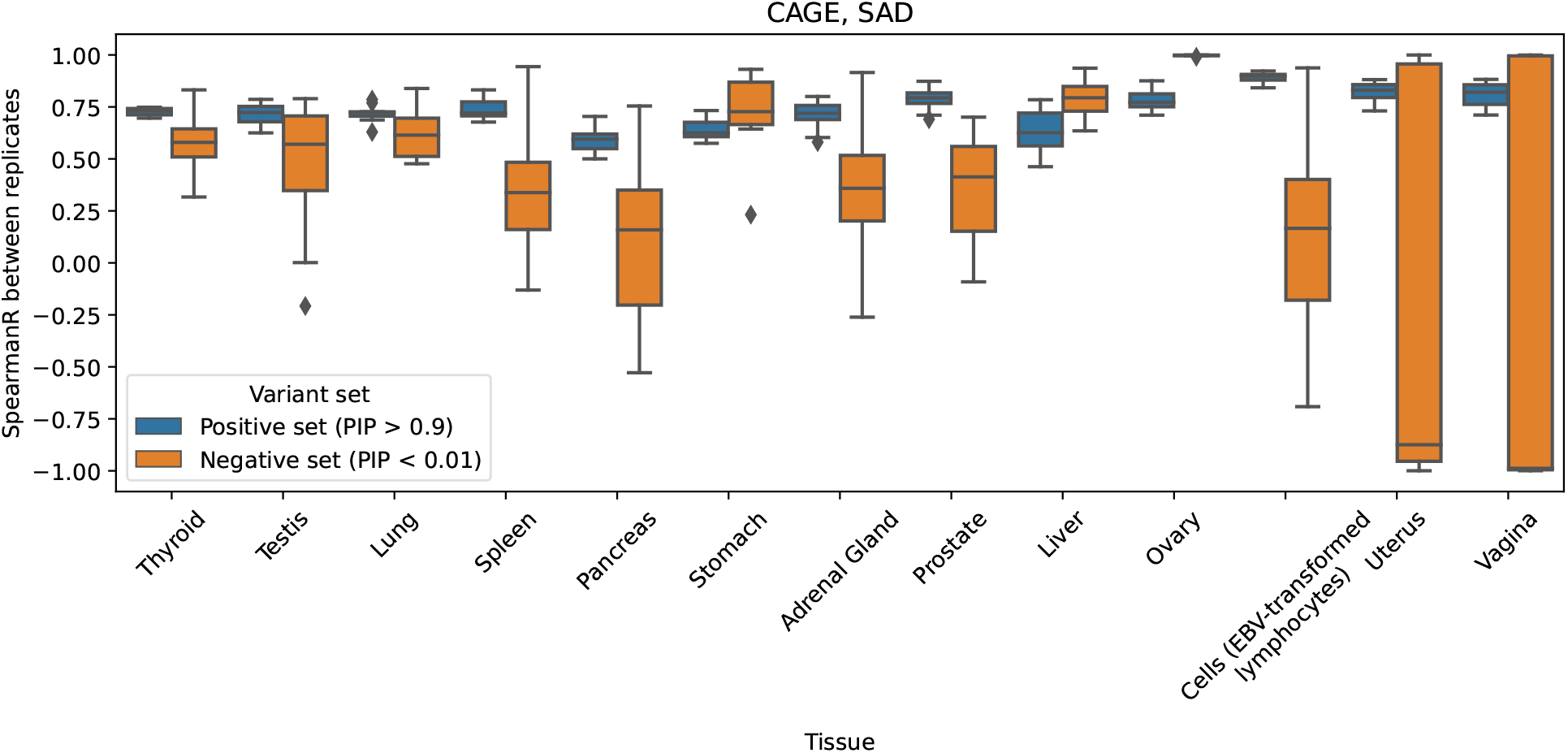
Replicates are more consistent in predicting the effects of fine-mapped eQTLs than the effects of variants in a matched negative set. For both sets of variants in each tissue, we calculate the Spearman correlation between predicted SAD (SNP Activity Difference) scores for every pair of replicates and show the distribution of correlations.

**Figure S5:**
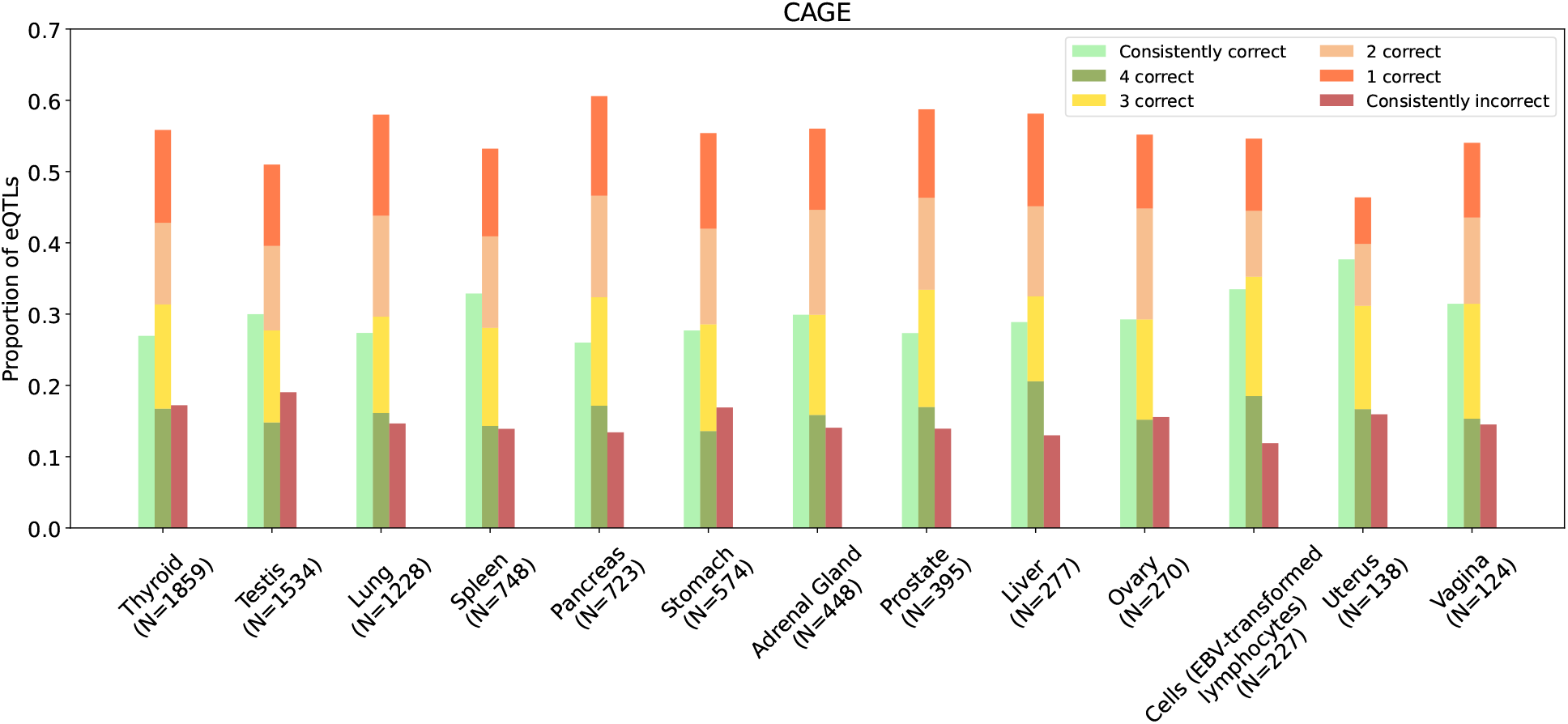
In all tissues, replicates have have inconsistent sign predictions for 50-60% of fine-mapped eQTLs when using the tissue-matched CAGE track. We stratify the plot in Fig. 3a by tissue and find similar proportions of consistently correct, inconsistent, and consistently incorrect sign predictions in all tissues.

**Figure S6:**
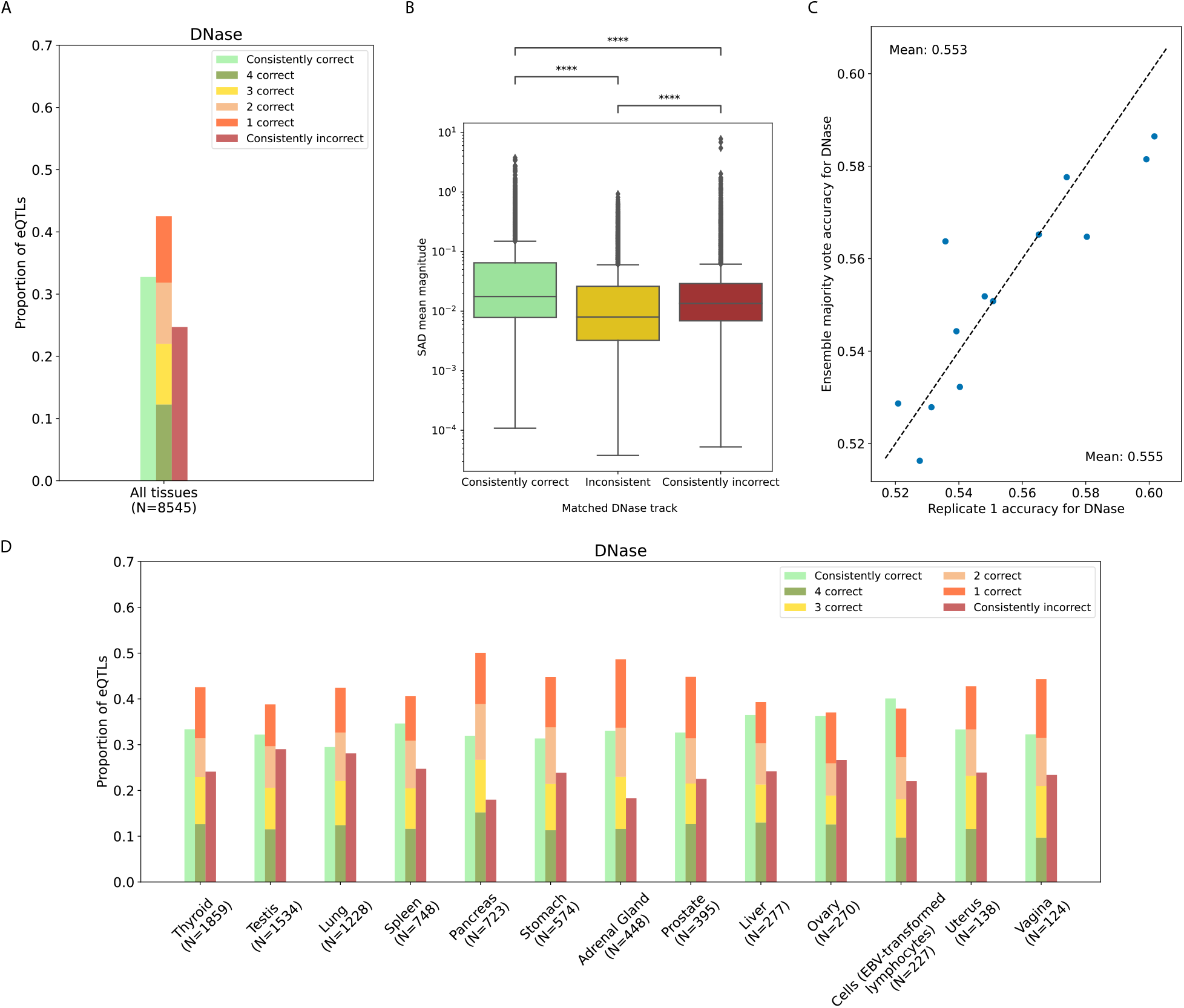
eQTL sign prediction for DNase has some inconsistency across replicates, but less than for CAGE. (a) We show the proportion of fine-mapped eQTLs with consistently correct, inconsistent, and consistently incorrect replicate predictions for eQTL effect sign. Pooled across tissues, about 32% of eQTLs are predicted consistently correctly, while about 25% are predicted consistently incorrectly. About 43% are inconsistently predicted. Of the consistently predicted eQTLs, about 65% are consistently correct, similar to the finding for CAGE. (b) eQTLs with inconsistent sign predictions across replicates tend to have smaller predicted effect sizes (mean of SAD score magnitude across replicates), for the tissue-matched DNase track. (c) A comparison of accuracy for eQTL sign prediction shows that the ensemble majority vote does not outperform a single replicate. Each point is the fine-mapped eQTL set of a different tissue. (d) We show the breakdown across tissues using matched DNase tracks. Across all tissues, about 40-50% of eQTLs have inconsistently predicted sign. About 30-40% eQTLs are predicted consistently correctly.

**Figure S7:**
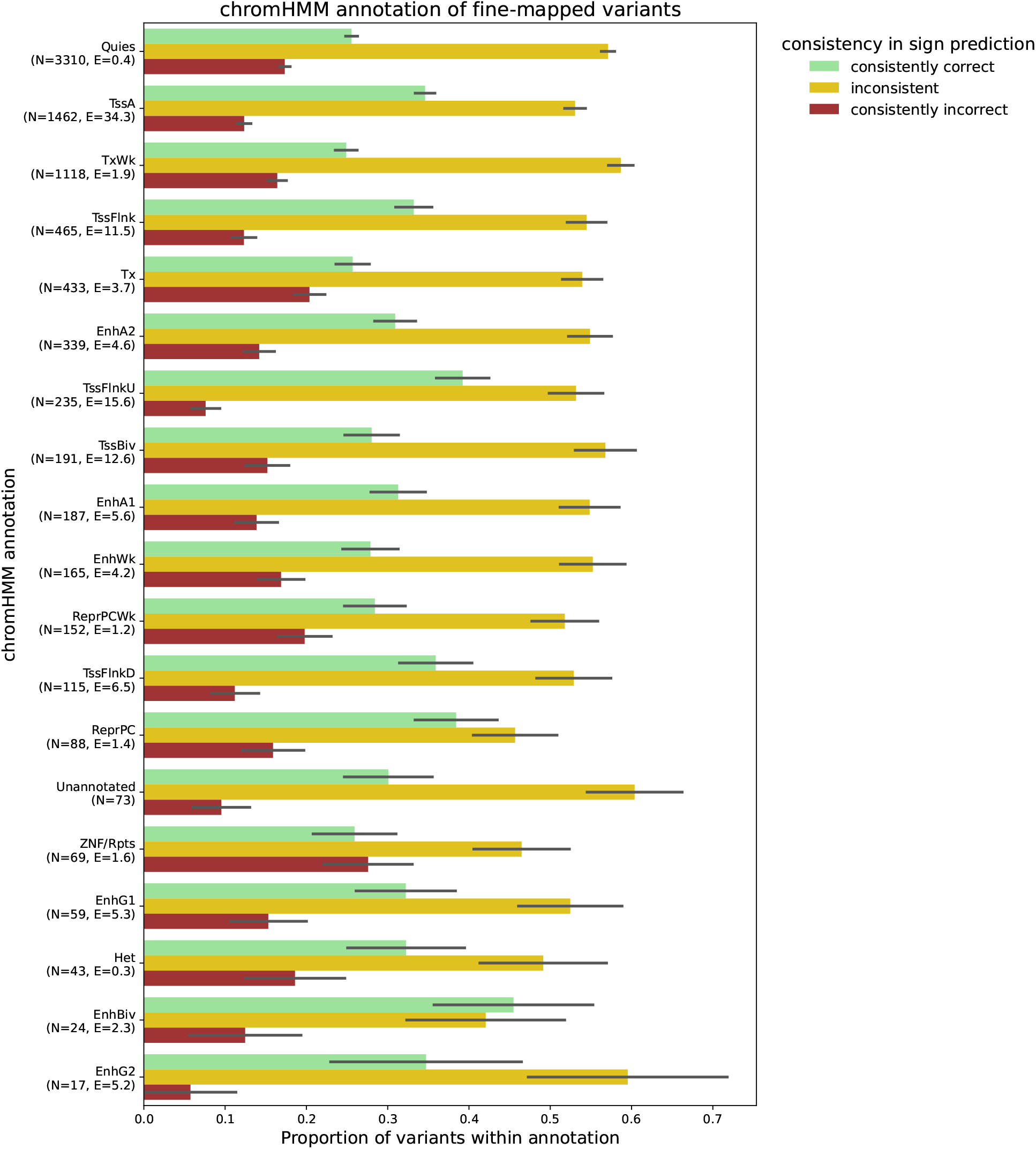
Replicate consistency in sign prediction of fine-mapped eQTLs segmented by chromHMM annotation. We labeled each fine-mapped variant with its tissue-matched chromHMM annotation or as “Unannotated” if it did not overlap a chromHMM annotation. For each annotation, we show the proportion of variants whose signs are predicted consistently correctly, inconsistently, or consistently incorrectly using the tissue-matched CAGE track. For all annotations, at least 40% of variants have inconsistent sign predictions. *N* denotes the number of fine-mapped variants overlapping that annotation, and *E* indicates the mean enrichment of fine-mapped variants in that annotation, averaged over the 13 examined tissues. Annotations are sorted by *N* . One standard deviation error bars are computed using 1000 bootstraps.

**Figure S8:**
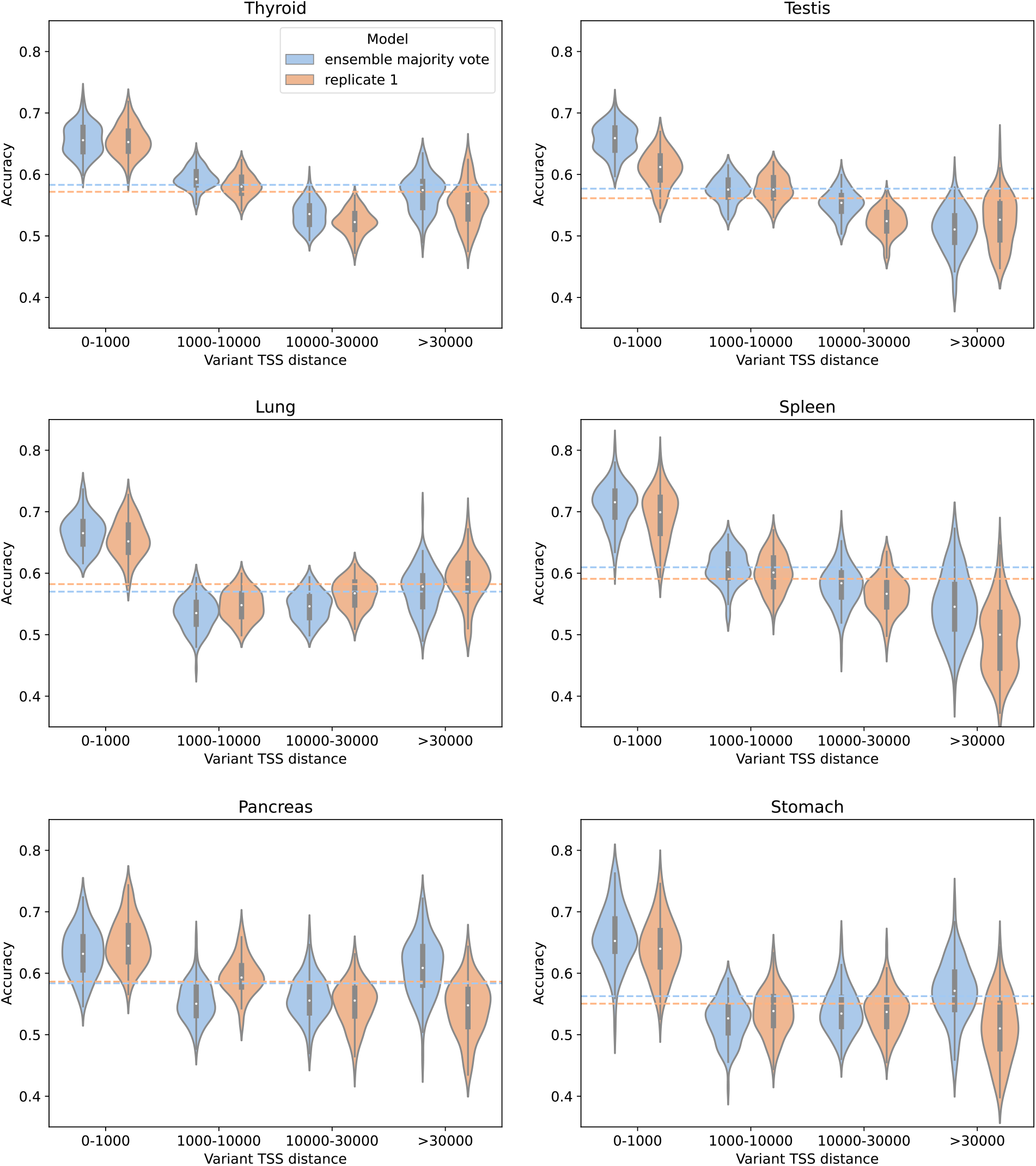
Ensemble majority vote for CAGE predictions often performs better on the subset of proximal variants. For the six tissues with the largest number of fine-mapped eQTLs, we stratify variants into TSS distance bins and compare eQTL sign prediction accuracy of a single replicate and the ensemble majority vote, using predictions from the tissue-matched CAGE track. Violinplots show the accuracy distributions over 100 bootstrap samples and dashed lines indicate the mean accuracies of the two models across all bins. In general, the ensemble majority vote has higher accuracy in the proximal TSS bin but not across all the bins. Across tissues, accuracy is highest for both models in the proximal TSS bin.

## Notes

### Competing Interest Statement

The authors have declared no competing interest.

